# PRISMOID: a comprehensive 3D structure database for post-translational modifications and mutations with functional impact

**DOI:** 10.1101/523308

**Authors:** Fuyi Li, Cunshuo Fan, Tatiana T. Marquez-Lago, André Leier, Jerico Revote, Cangzhi Jia, Yan Zhu, A. Ian Smith, Geoffrey I. Webb, Quanzhong Liu, Leyi Wei, Jian Li, Jiangning Song

**Author notes:** To whom correspondence should be addressed: (1) Fuyi Li, Biomedicine Discovery Institute, Department of Biochemistry and Molecular Biology, and Monash Centre of Data Science, Monash University, Melbourne, Victoria 3800, Australia.; (2) Leyi Wei, School of Computer Science and Technology, College of Intelligence and Computing, Tianjin University, Tianjin 300000, China.; (3) Quanzhong Liu, College of Information Engineering, Northwest A&F University, Yangling 712100, China.; (4) Jiangning Song, Biomedicine Discovery Institute, Department of Biochemistry and Molecular Biology, and Monash Centre of Data Science, Monash University, Melbourne, Victoria 3800, Australia.

## Abstract

Post-translational modifications (PTMs) play very important roles in various cell signalling pathways and biological process. Due to PTMs’ extremely important roles, many major PTMs have been thoroughly studied, while the functional and mechanical characterization of major PTMs is well-documented in several databases. However, most currently available databases mainly focus on protein sequences, while the real 3D structures of PTMs have been largely ignored. Therefore, studies of PTMs 3D structural signatures have been severely limited by the deficiency of the data. Here, we develop PRISMOID, a novel publicly available and free 3D structure database for a wide range of PTMs. PRISMOID represents an up-to-date and interactive online knowledge base with specific focus on 3D structural contexts of PTMs sites and mutations that occur on PTMs and in the close proximity of PTM sites with functional impact. The first version of PRISMOID encompasses 17,145 non-redundant modification sites on 3,919 related protein 3D structure entries pertaining to 37 different types of PTMs. Our entry web page is organized in a comprehensive manner, including detailed PTM annotation on the 3D structure and biological information in terms of mutations affecting PTMs, secondary structure features and per-residue solvent accessibility features of PTM sites, domain context, predicted natively disordered regions and sequence alignments. In addition, high-definition JavaScript packages are employed to enhance information visualization in PRISMOID. PRISMOID equips a variety of interactive and customizable search options and data browsing functions; these capabilities allow users to access data via keyword, ID, and advanced options combination search in an efficient and user-friendly way. A download page is also provided to enable users to download the SQL file, computational structural features, and PTM sites’ data. We anticipate PRISMOID will swiftly become an invaluable online resource, assisting both biologists and bioinformaticians to conduct experiments and develop applications supporting discovery efforts in the sequence-structural-functional relationship of PTMs and providing important insight into mutations and PTM sites interaction mechanisms. The PRISMOID database is freely accessible at http://prismoid.erc.monash.edu/. The database and web interface are implemented in MySQL, JSP, JavaScript, and HTML with all major browsers supported.

## 1. Introduction

The relatively modest number of human genes (~25,000) does not reflect the downstream proteomic complexity that is necessary for development and sustaining life. This essential complexity is achieved through a number of factors, such as protein post-translational modifications (PTMs), alternative splicing [1], intrinsic disorder [2, 3], etc. PTMs exhibit tremendous chemical diversity and dynamics, playing fundamental roles in various cell signalling pathways, fundamental cellular and biological processes. For example, phosphorylation, one of the most studied PTMs [4], is involved in regulation of protein degradation [5], signal transduction [6], protein-protein interaction [7], and signalling pathways [8]. Glycosylation, another important PTM, has been proved to play a crucial role in protein folding, trafficking [9-11], subcellular localization, and degradation [12, 13]. Other PTMs, such as lysine PTMs, are key for protein stability [14], transcription [15], cellular metabolism [16], DNA repair and replication [17] etc. Likewise, due to the fundamental importance of PTMs in cell biology, a number of aberrant PTM sites, which are often introduced by mutations, are highly relevant to various human diseases, including somatic cancers [18–20]. Altogether, it is critical to identify and characterize PTMs and their functional roles, which can in turn help elucidate the pathogenesis of PTMs’ related diseases.

A key step to understand mechanisms and functional roles of PTMs is to identify the sequences/structures of their natural substrates and the corresponding PTM sites. Given the fact that PTMs are strongly associated with mutations and diseases, significant efforts have been placed in using advanced high-throughput experimental techniques, such as mass spectrometry (MS) or MS-based techniques, to detect PTMs of proteins in proteomics data [21]. Therefore, large-scale modified proteomics data have been archived in recent years. To date, over 660 PTM types have been discovered (http://www.uniprot.org/docs/ptmlist.txt).

Several popular online databases have been developed, to collectively deposit and provide access to different types of experimentally validated PTMs data for public use. These databases include PhosphoSitePlus [22, 23], dbPTM [24, 25], SysPTM [26, 27], PLMD [28], Phospho.ELM [29], and UniProt [30]. PhosphoSitePlus was built by manually collecting and organizing data from various PTMs and model organisms, though primarily phosphorylation, ubiquitination and acetylation, from mainly human and mouse proteins. dbPTM has been maintained over 10 years, and integrates multiple PTMs information, including functional and structural knowledge of PTMs; dbPTM also stores drug binding data that shows associations to PTM sites. SysPTM contains both PTM data from public databases and MS experimental data from the literature. We note Pfam domains, KEGG pathways, and Gene Ontology information are also integrated into SysPTM. PLMD is a lysine-specific PTM database, and it encompasses 20 types of lysine PTMs in both eukaryotes and prokaryotes. Phospho.ELM is a phosphorylation-specific database, designed to provide general and kinase-specific phosphorylation sites collected from scientific literature and phosphoproteomic experiments. Lastly, UniProt is one of the largest and most regularly updated databases available, providing comprehensive annotations on protein sequences, functions and other relevant biological information. Currently, the UniProt database contains the information of over 660 types of PTMs.

These online resources collect protein sequences with experimentally validated post-translational modification sites, where the positions of PTM sites on sequences have also been documented. These publicly available online resources not only help biologists to conduct valuable experimental analyses, but also provide bioinformaticians with a wealth of resources to build novel prediction tools for studying sequences and functions of PTMs. All these, together, speed up PTMs’ research process.

However, there are few databases have been developed for systematically archiving and conveniently accessing 3D structural data of PTMs. PhosphoSitePlus provides 3D structure views in the current version, while this only focus on limited entries. PTM-SD [31] is a protein structure database focus on PTMs, which contains 11,677 entries, while PTM-SD only focus on the protein structure information of PTMs, other important information such as disorder information associated with the PTM site and PTM-associated disease mutations is not provided. In addition, PTM-SD only present the automatic produced figures of the protein 3D structure, the interactive visualization of protein 3D structures is not provided. The latest version of dbPTM provides the related PDB (Protein Data Bank) ID of PTM proteins, but the corresponding positions of PTM sites position on PDB structures are not available. Furthermore, dbPTM does not provide a download function for PTM related PDB information. Recently, Gao *et al*. [32] developed BioJavaMolFinder, a software designed to identify protein modifications stored in 3D structures within the PDB database. BioJavaMolFinder scanned PDB database and identified the proteins with modification, and provided the results on the PDB website. However, BioJavaMolFinder solely focuses on modification residues in PDBs, while the relationships among protein sequence-structure-function are not provided. More recently, Ledesma *et al*. developed YAAM for yeast amino acid modifications, which maps PTM sites in 680 protein 3D structures. On the whole, there is a generalized lack of user-friendly tools for both biologists and bioinformaticians to investigate the sequence-structural-function information of PTM proteins from existing protein sequence-based databases, limiting research efforts. For instance, although there are many bioinformatics prediction tools based on protein sequence data to investigate PTMs [4, 33–37], there are few prediction studies using PDB structures to build prediction models [38–40]. This is mainly due to the fact that the process of mapping from protein sequences to protein structures is quite time-consuming and operationally complex.

To bridge the existing knowledge gap, we developed a new web-based database, PRISMOID (<u>PR</u>ote<u>I</u>n <u>S</u>tructure <u>MO</u>d<u>I</u>fication <u>D</u>atabase) to provide a comprehensive annotation of the position of PTM sites in real 3D structures. We extracted experimentally validated PTMs information from several currently available protein sequence-based databases, and subsequently mapped the PTM sites to the corresponding 3D structures. We then filtrated the blast results and stored these results in PRISMOID. Aside, it has been reported there are 21% disease-related amino acid substitutions associated with missense single nucleotide variants (SNVs) are located in PTM sites [41]. Accordingly, we additionally extracted and collected PTM-associated disease and population mutations from ActiveDriverDB [41] database, so as to facilitate the investigation of structural properties of PTMs and to encourage construction of bioinformatics tools with structural features. To this effect, we also calculated secondary structure based features and residue accessibility features of PTM sites using DSSP [42] and NACCESS [43] software, respectively.

With all the above in consideration, PRISMOID provides users with the protein domain context, sequence-structural alignment and protein disorder information in each entry page. Moreover, to facilitate users’ access to data of interest, keyword/ID search, advanced combinatorial schemas search and “browse” functions are provided. PRISMOID also allows users to download the SQL file, the results files calculated from DSSP and NACCESS, and PTM annotation files in the download page.

## 2. Materials and Methods

### 2.1 PTM data collection

Figure 1 (A) shows the flowchart of PRISMOID steps, including data collection and processing. There are three major steps involved in entry collection and processing, described in what follows.

**Figure 1.**
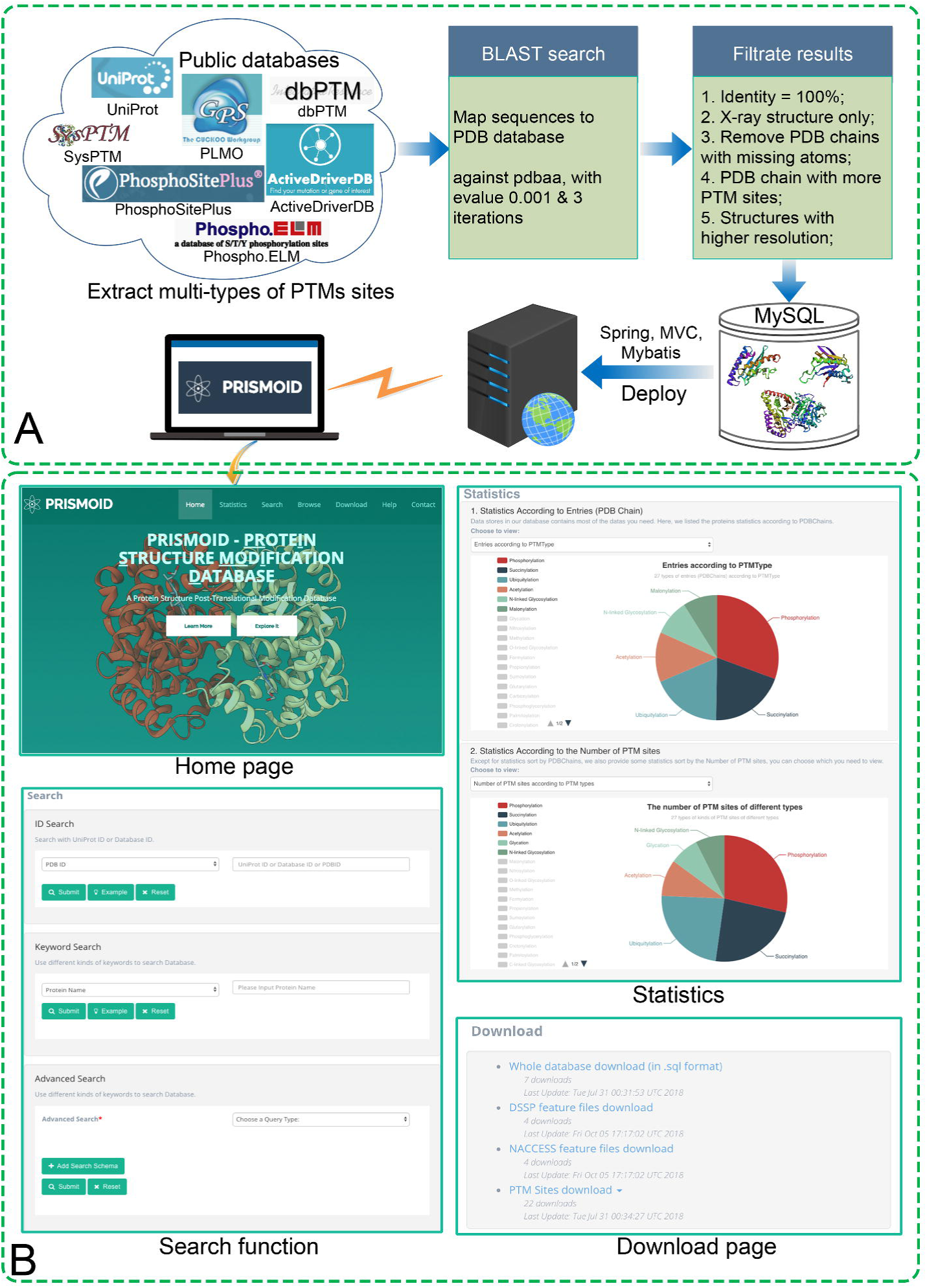
(**A**) PRISMOID data collection and processing flowchart; (**B**) Sample web interfaces. There are four major web pages in PRISMOID: “Home” page, “Search” page, “Statistics” page, and “Download” page.

First, we extracted, manually checked and integrated 37 types of PTM annotation data from six major online, public free available PTM databases, including PhosphoSitePlus [22], dbPTM [24], SysPTM [26], PLMD [28], Phospho.ELM [29], and UniProt [30]. We only considered and integrated experimentally validated PTM sites data for PRISMOID; accordingly, all PTM sites labelled as computational PTM annotations were discarded. For UniProt data, we only considered PTMs with ECO code ECO:0000269 (https://www.uniprot.org/help/evidences), which means the data is manually curated and involving published experimental data.

Secondly, all the remaining protein sequences with PTM sites were mapped to the PDB database [44] using PSI-BLAST [45] against PDB sequence database (pdbaa) with the e-value of 10^−3^ and three iterations.

Finally, the mapped PDB entries were selected according to the following criteria:

1. With 100% sequence identity; only PDB sequences aligned with the entry protein sequences with 100% sequence identity were retained.
2. X-ray resolution structures only; thus, nuclear magnetic resonance (NMR) and electron microscopy (EM) structures were excluded.
3. PDB chains with missing atoms were removed.
4. For each query blast sequence, the PDB chain with more PTM sites that can be aligned was selected;
5. For protein sequences with more than one mapped PDB structure, the structure with the highest resolution was selected.

After these five steps, a total number of 17,145 PTM sites on 3,919 PDB chains for 37 types of PTMs were extracted and stored into PRISMOID. The statistics of each type of PTM data are shown in **Table 1.** A discussion of the statistical analysis is included in Section 3.1.

### 2.2 PTM-associated Mutation collection

Single amino acid variants (SAVs) are the most abundant form of known genetic variations associated with human disease [46]. It is reported that about 60% of Mendelian diseases are caused by amino acid substitutions [47]. A potential effect of SAVs on protein function is the disruption of PTMs [48]. For example, the potential for phosphorylation sites to be affected by amino acid variants is high and there have been numerous examples of diseases associated naturally occurring variants that impact the phosphorylation status of proteins [49]. Thus, the mutations around PTMs sites are crucial in the study of health and disease. We have combined mutations associated with various diseases (including but not limited to cancer) and population variants affecting PTMs from ActiveDriverDB, which is a comprehensive collection of annotations of disease and population mutations contained in TCGA. The collected mutations were then mapped to the entries in PRISMOID. Please refer to Figure 2 (C) “Mutations affecting PTM sites” panel.

**Figure 2.**
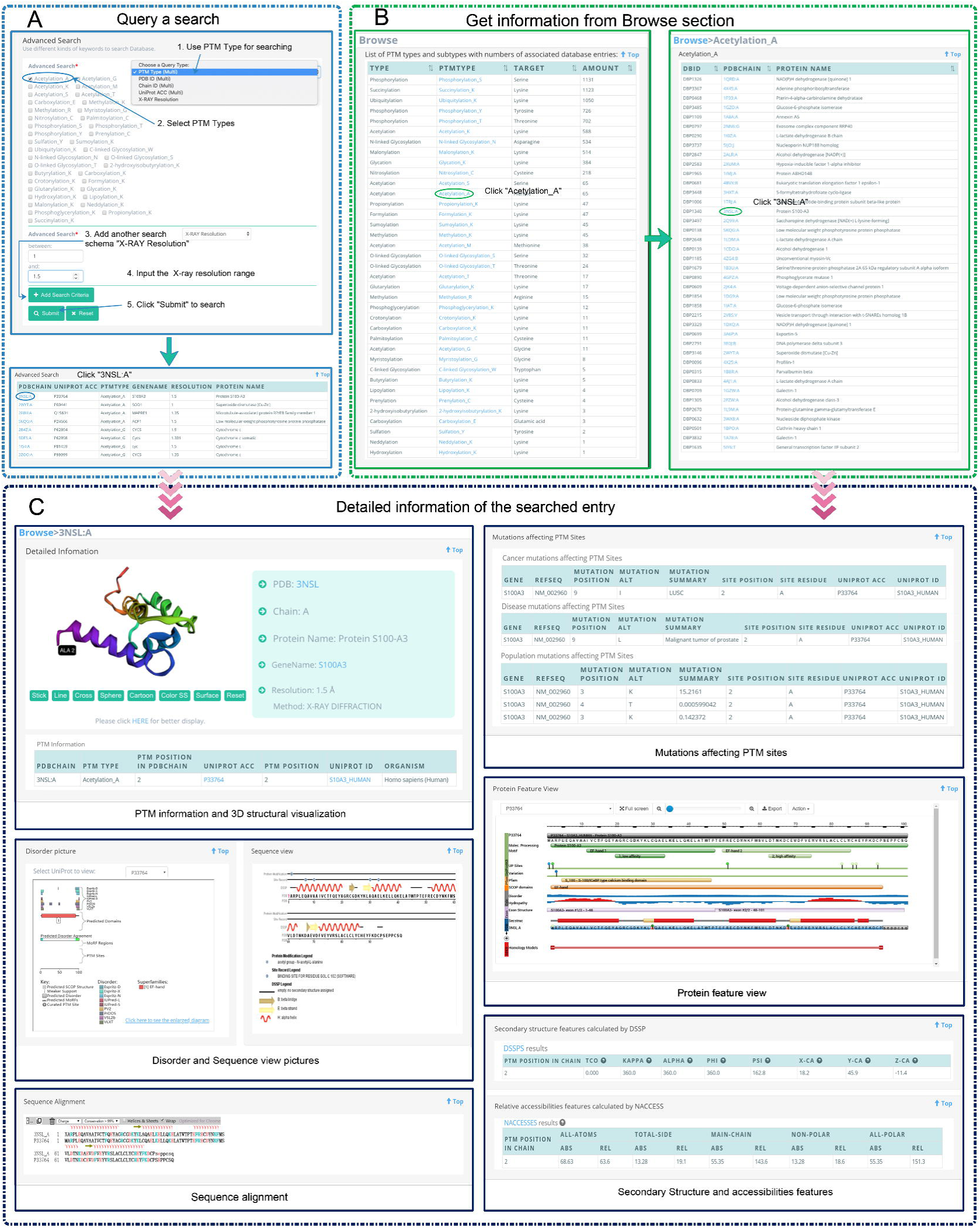
Two different methods for data accessing in PRISMOID. (**A**) An example showing how to use the advanced function of PRISMOID. Here, two different search schemas were combined, PTM types (Acetylation A is selected) and the range of X-ray resolution (between 1 and 1.5); (**B**) An example illustrates how to explore the PTM data through the Browse webpage (Acetylation A and PDB chain 3NSL:A is selected); (**C**) Typical search result webpage using the PDB chain 3NSL:A as an example. The entry webpage consists of six major sections showing detailed information: PTM information and 3D structural visualization, mutations affecting PTM sites information, protein feature view, secondary structure and accessibilities features, disorder and sequence view pictures and sequence alignment.

### 2.3 3D structure-based features calculation

Previous studies have shown that 3D structure-based features play an important role in improving the predictive performance of PTM sites [38, 50]. Therefore, we calculated two commonly used protein 3D structure features, secondary structure features and solvent accessible area features by using DSSP [42] and NACCESS [43], respectively. The result files can be downloaded from the download web page of PRISMOID ((http://prismoid.erc.monash.edu/pdb/download) please refer to Figure 1 (B)).

#### 2.3.1 Secondary structure features

Different types of features were extracted from the DSSP calculation result files for each PTM site, including secondary structure, bond angles (virtual bond angle, *KAPPA*), torsion angles (virtual torsion angle, *ALPHA;* IUPAC peptide backbone torsion angles, *PHI PSI*), atom coordinates (C_α_ atom coordinates, *X-CA Y-CA Z-CA*), and the number of water molecules (ACC) features. There are eight features for each PTM site shown on the corresponding entry page.

#### 2.3.2 Solvent accessible area (ASA) features

Since solvent accessible area (ASA) has been widely used in many bioinformatics fields, including hot spots identification [51], catalytic sites prediction [52], and glycosylation sites prediction [38], etc. Here, we employed the NACCESS program to calculate ASA features. The resulting file of NACCESS contains two types of ASA features, absolute ASA features and relative ASA features. Each type includes five different features: all-atoms, total-side, main-chain, non-polar, and all-polar ASA features. Thus, there are totally 10 ASA features for each PTM site. Please refer to the “Secondary structure and accessibilities features” panel of Figure 2 (C).

### 2.4 Database access and download functionality

To obtain a database with abundant data, it is essential to apply a good search algorithm to provide users with more efficient and convenient access. As shown in the “Search function” panel of Figure 1 (B), PRISMOID provides searching the entries in various ways, based on users’ favour. The entries can be searched through three different methods, including ID search, Keyword search, and Advanced search. The Advanced search method in PRISMOID allows multiple search schemas specific to different types of data combined, e.g. PTM types and PDB ID, or PDB Chain, UniProt Acc and X-Ray resolution range. The search function is described in detail in Section 3.1, and an example is given to illustrate the usage of the advanced search function.

PRISMOID also allows users to explore data embedded in the database through the “Browse” page. Figure 2 (B) illustrates an example of how to explore the data this way. When users click the “Browse” icon on the top page, the web page redirects them to “Browse” page, which displays a table with four columns including type (e.g. Acetylation), PTM type (e.g. Acetylation_A), target Amino acid (e.g. Alanine), and the number of entries (e.g. 65). The table lists all the PTM types stored in PRISMOID, as shown in the left sub-figure of Figure 2 (B). Then, when users click the PTM type (e.g. Acetylation_A), the web page redirects them to the browse page of this PTM. There users find a three-column table that includes database ID (e.g. DBP1340), PDB chain (e.g. 3NSL:A), and protein name (e.g. Protein S100-A3). Please refer to the right sub-figure of Figure 2 (B) for more details. Both the “Search” results page and “Browse” page allow results to be sorted, thereby enabling users to easily find the information they need, immediately.

PRISMOID also provides the download function to facilitate users download of PTMs data in PRISMOID. As shown in the “Download page” panel of Figure 1 (B), PRISMOID allows users to download the SQL file of the database, calculated DSSP feature files, calculated NACCESS feature files, and PTM sites data. These files can greatly facilitate bioinformaticians in their efforts to build machine-learning based bioinformatics methods for predicting PTMs sites straight from the protein structure level.

### 2.5 Implementation

The information stored in PRISMOID resides in a MySQL relational database consisting of 49 tables. All the routines in PRISMOID were written in the Java programming language. A highly user-friendly web front-end to the data was implemented using the jQuery framework. The basic operations of web interfaces are supported by JavaScript and jQuery, and the web page frame was built utilizing the bootstrap framework (https://getbootstrap.com/). Apache Tomcat 7 resides on a Linux server handles serving request of data from users, utilizing SSM (Spring, SpringMVC and MyBatis) framework for data searching, operation, web page hopping and viewing. The Linux server equipped with an 8-core CPU, 30 GB hard disk and 6 GB memory.

For the statistics page, the ECharts (https://ecomfe.github.io/echarts-doc/public/en/index.html) package [53] was used to achieve a user interactive pie charts display of statistical results. PRISMOID provides 11 different statistical results; users can view different statistics’ pie charts by selecting the different statistical methods from the drop-down menus. Additionally, when users move the mouse over a sector of a pie chart, the corresponding statistical results are displayed right where the cursor points. By default, the pie chart shows the statistical results of the top-five types. However, users can select the types to be displayed by clicking the legend on the left.

For the entry page, 3Dmol.js [54] was employed for protein 3D structure visualization. 3Dmol.js is a high-performance, pure object-oriented JavaScript library, enabling interactive visualization of protein 3D structures. We opted to use 3Dmod.js because it does not need to install browser plug-ins or Java. In PRISMOID, each entry is a specific PDB chain. Therefore, only the corresponding chain of the PDB structure is displayed, and the PTM sites on the chain are labelled at the corresponding positions. Please refer to the “PTM information and 3D structural visualization” panel in Figure 2 (C), where the PTM site is labelled at position 2 of the chain A as “ALA 2”. A Scalable Vector Graphics (SVG) library, the Protein Feature view (http://andreasprlic.github.io/proteinfeatureview/), was applied to show variation data, Pfam domains, functional sites and some basic structural information. This visualization library enables users to quickly identify PTMs that lie within functional domains and to visualize the topology of the PTMs. The “Protein feature view” panel in Figure 2 (C) illustrates this function. The RCSB-sequence viewer plugin (https://github.com/bioiava/rcsb-sequenceviewer) was also employed to generate the sequence view picture of the PDB chain. The sequence view shows the protein modification generated by BioJavaMolFinder, PDB chain sequence and secondary structures from DSSP. In addition, we also extracted the protein disorder figures from the disordered protein predictions database D2P2 [55]. An example of the sequence view and protein disorder pictures are shown in the “Disorder and Sequence view pictures” panel in Figure 2 (C). Lastly, Alignment-To-HTML [56] was used for interactive visualization for the sequence alignments between the PDB chain sequence and the corresponding UniProt sequences.

## 3. Results and Discussion

### 3.1 Search functionality

As we discussed in section 2.4, PRISMOID provides several search algorithms, enabling users to get easy, comprehensive access to data. Here we introduce the search function of PRISMOID in detail, and present a specific example to show how to use the advanced search function.

PRISMOID data can be searched by three different search methods. The first method is the ID search, where the users can search PRISMOID entries by using a PDB ID (e.g. ‘3NSL’), UniProt ACC (e.g. ‘P33764’) or Database ID (e.g. ‘DBP1340’). The second method is the keyword search, where users can search entries by entering a protein name or gene name. For these two search methods, a corresponding ‘Example’ button is provided, to guide users to conduct a search. With the click of the ‘Example’ button, users can automatically get an example for a search with ID/keyword. The third method is the Advanced Search. There are five different query schemas in the Advanced search field, including (i) PTM types, (ii) PDB IDs, (iii) Chain IDs, (iv) UniProt ACCs; and (v) range of X-Ray resolution. Users can search with individual schema or combinations of multiple schemas by selecting the search schema in the dropdown list of the ‘Advanced search’ menu and using the “Add Search Schema” button to search with multiple search schemas. Figure 2 (A) illustrates an example using the combination of PTM types (“Acetylation_A” was selected) and X-ray resolution range (1 to 1.5 were filled) to conduct the search.

Searched results will be shown in a table on a separate webpage. The table will contain six columns, showing basic information of searched queries, including PDB ID, chain ID, PTM positions in PDB chain, the UniProt accession number (Acc), PTM positions in UniProt sequence, organism, PTM type, gene name, PDB resolution, and protein name (as shown in Figure 2 (A)). Moreover, detailed information of each entry can be browsed in a new webpage by clicking the PDB Chain in the result table. For each entry, there are eight major types of information provided, including protein 3D structure visualization, PTM information, mutations affecting PTM sites information, secondary structure based features calculated by DSSP, relative accessibilities features calculated by NACCESS, protein feature view, sequence alignment, disorder picture, and protein sequence view. To provide a demonstration of the annotation information for each entry in PRISMOID, ‘PDBChain=3NSL:A’ (PRISMOID ID=‘DBP1340’) was used as an example query. The resulting page with annotation information is shown in Figure 2 (C).

### 3.2 Database contents

With the data contained in PRISMOID, we conducted a statistical analysis of the PTM information distribution across the top eight species with the largest number of entries/PTM sites. Two bar charts (Figure 3 (A) and (B)) show statistical results of the eight species with the largest number of entries (PDB chain) and PTM sites, respectively. The most abundant source with 11,560 PTM sites on 2,171 PDB chains was Human, followed by *E. coli* with 2,246 PTM sites on 549 entries, Yeast with 922 PTM sites on 275 entries, and Mouse with 877 PTM sites on 219 entries.

**Figure 3.**
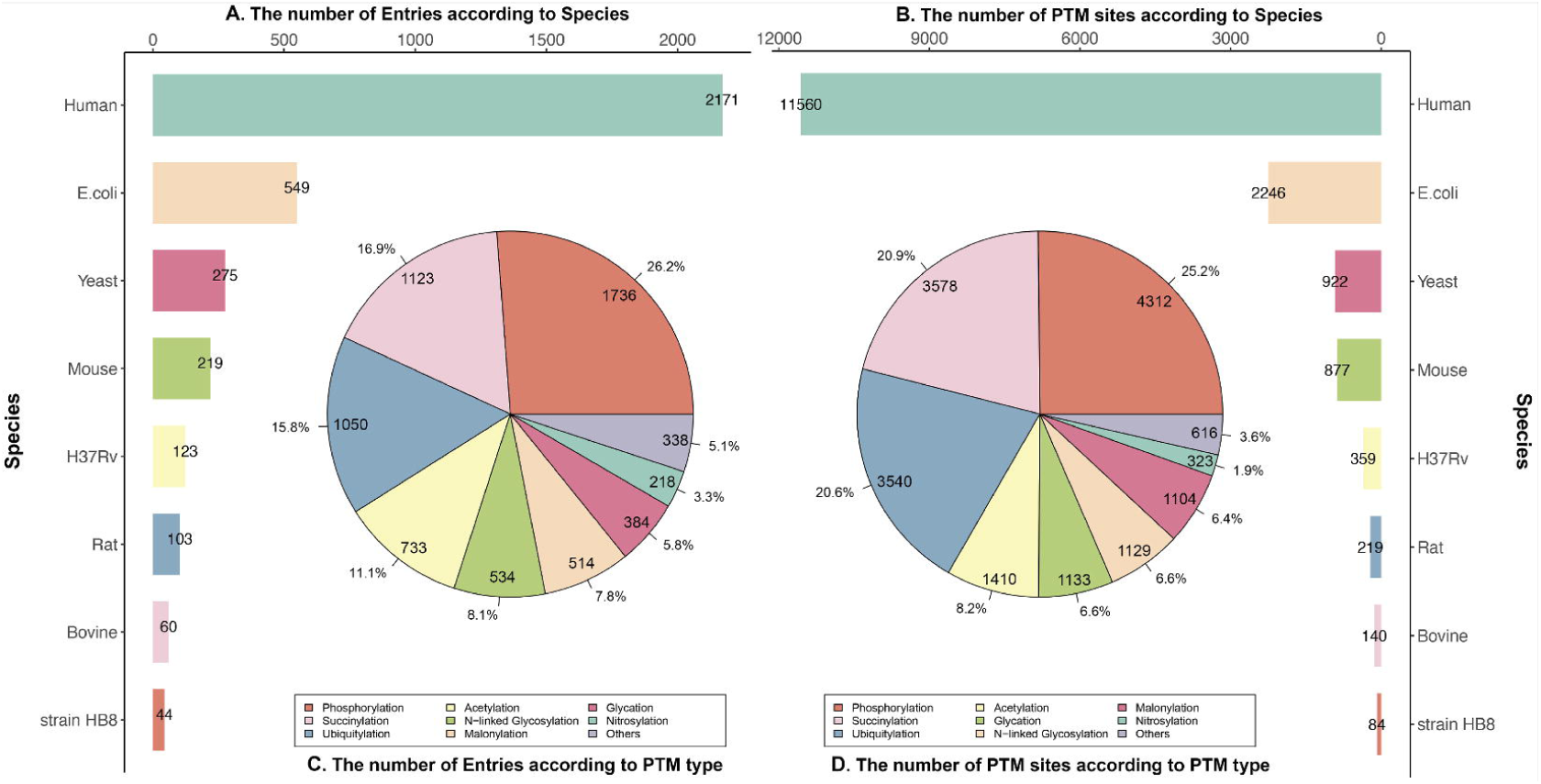
Statistical analysis results in terms of species and PTM types. (**A**) Statistical results of the entries (PDB chains) from the eight largest species, including *Homo sapiens, Escherichia coli, Saccharomyces cerevisiae, Mus musculus, Mycobacterium tuberculosis, Rattus norvegicus, Bos taurus*, and *Thermus thermophiles;* (**B**) Statistical analysis of the PTM sites from the eight largest species; (**C**) Distributions of eight largest PTM types in terms of the number of PTM entries; and (**D**) Distributions of eight largest PTM types in terms of the number of PTM sites.

Two pie charts in Figure 3 (C and D) indicate the distribution of entries and PTM sites of the eight largest number of PTM types, respectively. In general, the top eight largest types of PTM are the same, but the order is not exactly the same, in terms of the number of PTM proteins (entries) and the number of PTM sites. For example, the top two PTM types in terms of the number of proteins and PTM sites are the same, i.e. Phosphorylation and Acetylation respectively. There are 1736 phosphorylated proteins with 4312 phosphorylation sites and 1123 acetylated proteins with 3578 acetylation sites. The third largest type in terms of the PTM proteins number is Glycation (1050), while the third largest type in terms of the PTM sites number is Malonylation (3540).

The bar chart in Figure 4 illustrates the number of PTM sites associated with mutations. There are three types of mutations, including somatic cancer mutations, inherited disease mutations, and population mutations. **Table 2 and Supplementary Table S1** list the cancer types and disease types of somatic cancer mutations and disease mutations, respectively.

**Figure 4.**
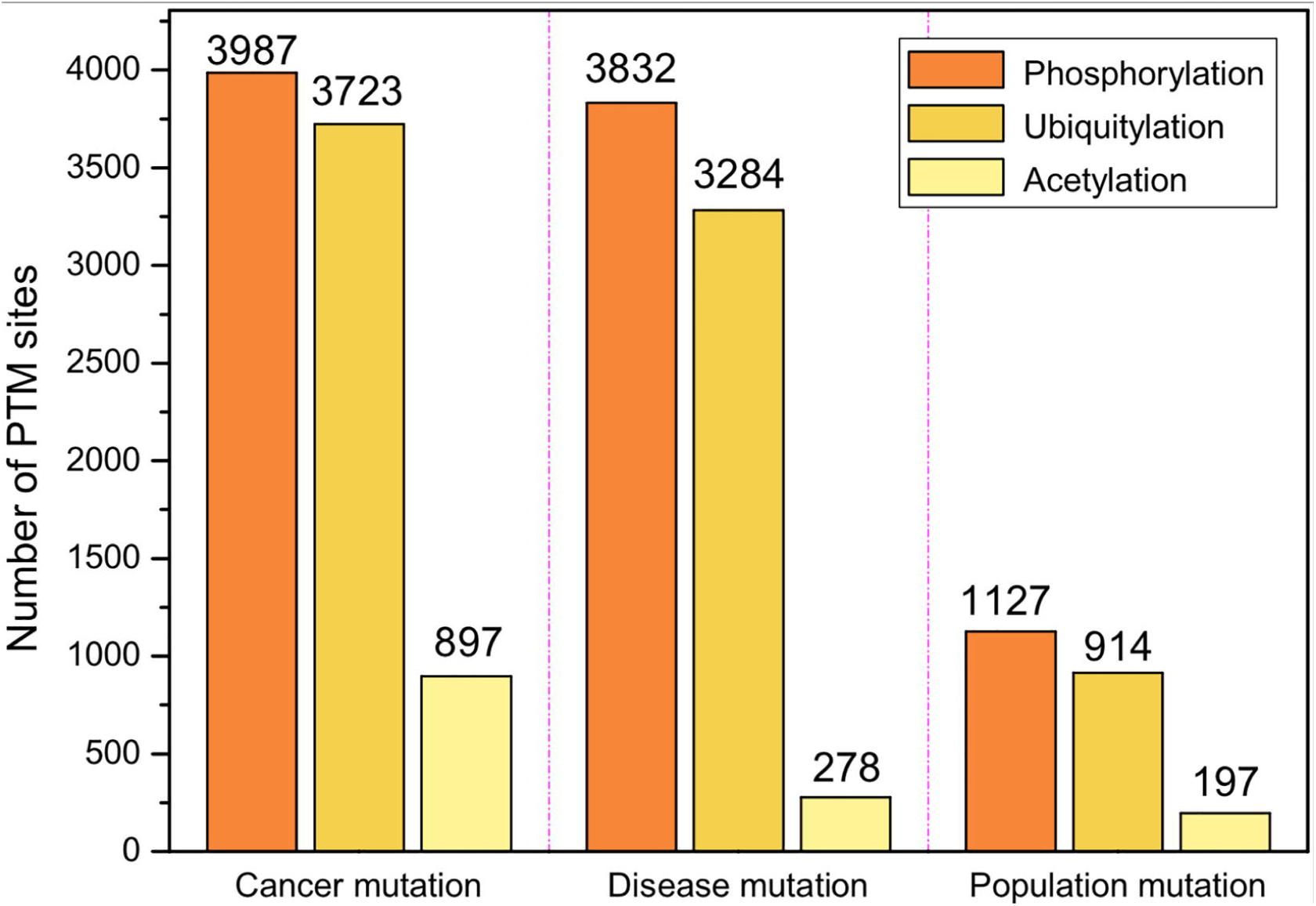
The top three PTMs in terms of the number of PTM sites associated with mutations. There are three types of mutations, including somatic cancer mutations, inherited disease mutations, and population mutations.

The top three types of PTMs associated with three types of mutations are the same. They are: (i) Phosphorylation with 3987 cancer mutations, 3832 disease mutations, and 1127 population mutations; (ii) Ubiquitylation with 3723 cancer mutations, 3284 disease mutations, and 914 acetylation mutations; and (iii) Acetylation with 897 cancer mutations, 278 disease mutations and 197 population mutations.

## 4. Conclusions

PRISMOID is a freely available web-based resource providing 3D structure information of protein post-translational modification sites. PRISMOID retrieves comprehensive PTM information from six major protein sequence-based PTM databases. The collected protein sequences with PTM sites were then mapped to protein 3D structures. PRISMOID not only includes PTM sites data but also integrates and annotates the disease/cancer/population mutation affecting the PTM sites. Other information, such as protein secondary structure properties, solvent accessibility area features, protein disorder region, and protein domain context, etc., were also collected and annotated in PRISMOID. The annotations of 3D structural features of PTM sites in PRISMOID can effectively guide the design of experimental efforts such as crystallization and mutagenesis to interrogate the functional roles of PTM sites.

PRISMOID combines the strength of Java, Spring, SpringMVC, MyBatis, and JavaScript to achieve user-friendly interface design and efficient data query/display. Multiple data querying methods and data browsing pages enabled in PRISMOID can be explored, with a number of options. PRISMOID is also equipped with several bioinformatics plug-ins such as 3Dmol.js, Protein Feature view, and RCSB-sequence viewer to better visualize the comprehensive annotation data contained in PRISMOID. As new datasets accumulate rapidly, as well as computational resources, it is essential to maintain the database current. Thus, we plan to develop a website background management system to facilitate timely and automatized updates of the database at least once every six months. In the future, we will incorporate additional information such as PTM crosstalk patterns into PRISMOID. A straightforward mechanism of two PTM sites to be associated in a cross-talk pattern is based on their spatial proximity [57, 58]. Accordingly, the 3D structural information can be used to calculate the spatial distance between any two PTM sites and help identify potential PTM crosstalk pairs. In addition, it has been reported that some PTM sites tend to be located in disordered regions [59]. In view of this, PRISMOID has included the predicted disorder annotations for all related entries by providing the link to the D2P2 database [55]. In an upgraded version of PRISMOID in the future, we aim to collect the reliably annotated disordered region information from multiple disordered region databases (including D2P2 and others) and map the disordered region to the PDB structures of proteins with PTM sites to enable users to perform an in-depth analysis of the potential association between PTM sites and disorder regions. Taken altogether, we anticipate that PRISMOID will become a useful resource for both experimental and computational analyses of protein post-translational modifications.

## Supporting information

The statistics of each type of PTM data are shown in Table S1

Table 2 and Supplementary Table S1 list the cancer types and disease types of somatic cancer mutations and disease mutations, respectively.

Table 2 and Supplementary Table S1 list the cancer types and disease types of somatic cancer mutations and disease mutations, respectively.

## Author Biography

**Fuyi Li** received his BEng and MEng degrees in Software Engineering from Northwest A&F University, China. He is currently a PhD candidate in the Department of Biochemistry and Molecular Biology and the Infection and Immunity Program, Biomedicine Discovery Institute, Monash University, Australia. His research interests are bioinformatics, computational biology, machine learning, and data mining.

**Cunshuo Fan** is an undergraduate student in the College of Information Engineering, Northwest A&F University, China. His research interests are bioinformatics, machine learning, data mining, and software engineering.

**Tatiana T. Marquez-Lago** is an associate professor in the Department of Genetics and the Department of Cell, Developmental and Integrative Biology, University of Alabama at Birmingham (UAB) School of Medicine, USA. Her research interests include multiscale modelling and simulations, artificial intelligence, bioengineering and systems biomedicine. Her interdisciplinary lab studies stochastic gene expression, chromatin organization, antibiotic resistance in bacteria, and host-microbiota interactions in complex diseases.

**André Leier** is currently an assistant professor in the Department of Genetics, University of Alabama at Birmingham (UAB) School of Medicine, USA. He is also an associate scientist in the UAB Comprehensive Cancer Centre. He received his Ph.D. in Computer Science (Dr. rer. nat.), University of Dortmund, Germany. He conducted postdoctoral research at Memorial University of Newfoundland, Canada, The University of Queensland, Australia and ETH Zürich, Switzerland. His research interests are in biomedical informatics and computational and systems biomedicine.

**Jerico Revote** received his bachelor degree in Computer Science from RMIT University, Melbourne, Australia. He is working as a research engineer in the Monash eResearch Centre at Monash University, Australia. He is also currently a part-time PhD student in the Department of Biochemistry and Molecular Biology and Biomedicine Discovery Institute, Monash University. His research interests are machine learning, data mining, artificial intelligence, software development, and scalable applications.

**Cangzhi Jia** is an assistant professor in the College of Science, Dalian Maritime University. She obtained her Ph.D. degree in school of mathematical sciences from Dalian University of Technology in 2007. Her major research interests include mathematical modelling in bioinformatics and machine learning.

**Yan Zhu** is a research fellow in Monash Biomedicine Discovery Institute and Department of Microbiology, Monash University, Australia. He is currently an NHMRC Principal Research Fellow. His research interests include the pharmacology of polymyxins and the discovery of novel, safer polymyxins.

**A. Ian Smith** completed his PhD at Prince Henrys Institute Melbourne and Monash University, Australia. He is currently the Vice-Provost (Research & Research Infrastructure) at Monash University. He is also a Professorial Fellow in the Department of Biochemistry and Molecular Biology at Monash University, where he runs his research group. His research applies proteomic technologies to study the proteases involved in the generation and metabolism of peptide regulators involved in both brain and cardiovascular function.

**Geoffrey I. Webb** received his PhD degree in 1987 from La Trobe University. He is the director of the Monash Centre for Data Science and Professor in Faculty of Information Technology at Monash University, Australia. He is a leading data scientist and have been the Program Committee Chair of two leading Data Mining conferences, ACM SIGKDD and IEEE ICDM. His research interests include machine learning, data mining, computational biology, and user modelling.

**Quanzhong Liu** is an associate professor in the College Information Engineering, Northwest A&F University, Yangling, China. He received his PhD in Agricultural Electrisation & Automatization from Northwest A&F University, China. His research interests include machine learning, data mining, big data and bioinformatics.

**Leyi Wei** received his PhD degree in Computer Science from Xiamen University, China. He is currently an assistant professor at the School of Computer Science and Technology, Tianjin University, China. His research interests include machine learning and their applications to bioinformatics.

**Jian Li** is a Professor and group leader in Monash Biomedicine Discovery Institute and Department of Microbiology, Monash University, Australia. He is a Web of Science 2015-2017 Highly Cited Researcher in Pharmacology & Toxicology. He is currently an NHMRC Principal Research Fellow. His research interests include the pharmacology of polymyxins and the discovery of novel, safer polymyxins.

**Jiangning Song** is an Associate Professor and group leader in the Monash Biomedicine Discovery Institute, Monash University, Melbourne, Australia. He is also affiliated with the Monash Centre for Data Science, Faculty of Information Technology, Monash University. His research interests include bioinformatics, computational biology, machine learning, data mining, and pattern recognition.

## Acknowledgements

This work was supported by grants from the National Health and Medical Research Council of Australia (NHMRC) (1144652 and 1127948), the Australian Research Council (ARC) (LP110200333 and DP120104460), the National Institute of Allergy and Infectious Diseases of the National Institutes of Health (R01 AI111965), and a Monash Major Inter-Disciplinary Research (IDR) Grant. Q.L. is supported by grants from the Key Research and Development Program of Shaanxi Province, China (2017GY-197). TML and AL’s work was supported in part by the Informatics Institute of the School of Medicine at UAB. J.L. is an NHMRC Principal Research Fellow.

## Key Points

- PRISMOID is a novel freely available web-based resource providing 3D structure information of a wide range of protein post-translational modification sites.
- The first version of PRISMOID encompasses 17,145 non-redundant modification sites on 3,919 related protein 3D structure entries pertaining to 37 different types of PTMs.
- The entry web page of PRISMOID is organized in a comprehensive manner, including detailed PTM annotation on the 3D structure and biological information in terms of mutations affecting PTMs, secondary structure features and per-residue solvent accessibility features of PTM sites, domain context, predicted natively disordered regions and sequence alignments.
- PRISMOID equips a variety of interactive and customizable search options and data browsing functions; these capabilities allow users to access data via keyword, ID, and advanced options combination search in an efficient and user-friendly way. A download page is also provided to enable users to download the SQL file, computational structural features, and PTM sites’ data.

## References

1. Pan Q, Shai O, Lee LJ et al. Deep surveying of alternative splicing complexity in the human transcriptome by high-throughput sequencing, Nat Genet 2008;40:1413-1415.

2. Peng ZL, Yan J, Fan X et al. Exceptionally abundant exceptions: comprehensive characterization of intrinsic disorder in all domains of life, Cellular and Molecular Life Sciences 2015;72:137-151.

3. Meng F, Uversky VN, Kurgan L. Comprehensive review of methods for prediction of intrinsic disorder and its molecular functions, Cellular and Molecular Life Sciences 2017;74:3069-3090.

4. Li F, Li C, Marquez-Lago TT et al. Quokka: a comprehensive tool for rapid and accurate prediction of kinase family-specific phosphorylation sites in the human proteome, Bioinformatics 2018:bty522–bty522.

5. Swaney DL, Beltrao P, Starita L et al. Global analysis of phosphorylation and ubiquitylation crosstalk in protein degradation, Nat Methods 2013;10:676–682.

6. McCubrey JA, May WS, Duronio V et al. Serine/threonine phosphorylation in cytokine signal transduction, Leukemia 2000;14:9–21.

7. Nishi H, Hashimoto K, Panchenko AR. Phosphorylation in protein-protein binding: effect on stability and function, Structure 2011;19:1807–1815.

8. Duan G, Walther D. The roles of post-translational modifications in the context of protein interaction networks, PLoS Comput Biol 2015;11:e1004049.

9. Moharir A, Peck SH, Budden T et al. The role of N-glycosylation in folding, trafficking, and functionality of lysosomal protein CLN5, Plos One 2013;8:e74299.

10. Marino K, Bones J, Kattla JJ et al. A systematic approach to protein glycosylation analysis: a path through the maze, Nat Chem Biol 2010;6:713–723.

11. Li F, Zhang Y, Purcell AW et al. Positive-unlabelled learning of glycosylation sites in the human proteome, BMC Bioinformatics 2019;20:112.

12. Dwek RA. Biological importance of glycosylation, Characterization Of Biotechnology Pharmaceutical Products 1998;96:43–47.

13. von der Lieth CW, Bohne-Lang A, Lohmann KK et al. Bioinformatics for glycomics: status, methods, requirements and perspectives, Briefings In Bioinformatics 2004;5:164–178.

14. Polevoda B, Sherman F. The diversity of acetylated proteins, Genome Biol 2002;3:reviews0006.

15. Glozak MA, Sengupta N, Zhang X et al. Acetylation and deacetylation of non-histone proteins, Gene 2005;363:15–23.

16. Zhao S, Xu W, Jiang W et al. Regulation of cellular metabolism by protein lysine acetylation, Science 2010;327:1000–1004.

17. Pickart CM. Ubiquitin enters the new millennium, Mol Cell 2001;8:499–504.

18. Karaca M, Liu Y, Zhang Z et al. Mutation of androgen receptor N-terminal phosphorylation site Tyr-267 leads to inhibition of nuclear translocation and DNA binding, PLoS One 2015;10:e0126270.

19. Fleuren EDG, Zhang LX, Wu JM et al. The kinome ‘at large’ in cancer, Nature Reviews Cancer 2016;16:83–98.

20. Pinho SS, Reis CA. Glycosylation in cancer: mechanisms and clinical implications, Nat Rev Cancer 2015;15:540–555.

21. Medzihradszky KF. Peptide sequence analysis, Methods Enzymol 2005;402:209–244.

22. Hornbeck PV, Zhang B, Murray B et al. PhosphoSitePlus, 2014: mutations, PTMs and recalibrations, Nucleic Acids Res 2015;43:D512–520.

23. Hornbeck PV, Kornhauser JM, Tkachev S et al. PhosphoSitePlus: a comprehensive resource for investigating the structure and function of experimentally determined post-translational modifications in man and mouse, Nucleic Acids Res 2012;40:D261–270.

24. Huang KY, Su MG, Kao HJ et al. dbPTM 2016: 10-year anniversary of a resource for post-translational modification of proteins, Nucleic Acids Res 2016;44:D435–446.

25. Lee TY, Huang HD, Hung JH et al. dbPTM: an information repository of protein post-translational modification, Nucleic Acids Res 2006;34:D622–627.

26. Li J, Jia J, Li H et al. SysPTM 2.0: an updated systematic resource for post-translational modification, Database (Oxford) 2014;2014:bau025.

27. Li H, Xing X, Ding G et al. SysPTM: a systematic resource for proteomic research on post-translational modifications, Mol Cell Proteomics 2009;8:1839–1849.

28. Xu H, Zhou J, Lin S et al. PLMD: An updated data resource of protein lysine modifications, J Genet Genomics 2017;44:243–250.

29. Dinkel H, Chica C, Via A et al. Phospho.ELM: a database of phosphorylation sites--update 2011, Nucleic Acids Res 2011;39:D261–267.

30. UniProt Consortium T. UniProt: the universal protein knowledgebase, Nucleic Acids Res 2018;46:2699.

31. Craveur P, Rebehmed J, de Brevern AG. PTM-SD: a database of structurally resolved and annotated posttranslational modifications in proteins, Database (Oxford) 2014;2014.

32. Gao J, Prlic A, Bi C et al. BioJava-ModFinder: identification of protein modifications in 3D structures from the Protein Data Bank, Bioinformatics 2017;33:2047–2049.

33. Li F, Li C, Wang M et al. GlycoMine: a machine learning-based approach for predicting N-, C- and O-linked glycosylation in the human proteome, Bioinformatics 2015;31:1411–1419.

34. Jia CZ, Zuo Y, Zou Q. O-GlcNAcPRED-II: an integrated classification algorithm for identifying O-GlcNAcylation sites based on fuzzy undersampling and a K-means PCA oversampling technique, Bioinformatics 2018;34:2029–2036.

35. Xue Y, Ren J, Gao X et al. GPS 2.0, a tool to predict kinase-specific phosphorylation sites in hierarchy, Mol Cell Proteomics 2008;7:1598–1608.

36. Chen Z, Zhou Y, Zhang Z et al. Towards more accurate prediction of ubiquitination sites: a comprehensive review of current methods, tools and features, Brief Bioinform 2015;16:640–657.

37. Taherzadeh G, Yang YD, Xu HD et al. Predicting lysine-malonylation sites of proteins using sequence and predicted structural features, Journal of Computational Chemistry 2018;39:1757–1763.

38. Li F, Li C, Revote J et al. GlycoMine(struct): a new bioinformatics tool for highly accurate mapping of the human N-linked and O-linked glycoproteomes by incorporating structural features, Sci Rep 2016;6:34595.

39. Chuang GY, Boyington JC, Joyce MG et al. Computational prediction of N-linked glycosylation incorporating structural properties and patterns, Bioinformatics 2012;28:2249–2255.

40. Durek P, Schudoma C, Weckwerth W et al. Detection and characterization of 3D-signature phosphorylation site motifs and their contribution towards improved phosphorylation site prediction in proteins, BMC Bioinformatics 2009;10.

41. Krassowski M, Paczkowska M, Cullion K et al. ActiveDriverDB: human disease mutations and genome variation in post-translational modification sites of proteins, Nucleic Acids Res 2018;46:D901–D910.

42. Joosten RP, te Beek TA, Krieger E et al. A series of PDB related databases for everyday needs, Nucleic Acids Res 2011;39:D411–419.

43. Hubbard S. NACCESS: program for calculating accessibilities, Department of Biochemistry and Molecular Biology, University College of London 1992.

44. Berman HM, Westbrook J, Feng Z et al. The Protein Data Bank, Nucleic Acids Res 2000;28:235–242.

45. Altschul SF, Madden TL, Schaffer AA et al. Gapped BLAST and PSI-BLAST: a new generation of protein database search programs, Nucleic Acids Res 1997;25:3389–3402.

46. Yip YL, Scheib H, Diemand AV et al. The Swiss-Prot variant page and the ModSNP database: a resource for sequence and structure information on human protein variants, Hum Mutat 2004;23:464–470.

47. Botstein D, Risch N. Discovering genotypes underlying human phenotypes: past successes for mendelian disease, future approaches for complex disease, Nat Genet 2003;33 Suppl:228–237.

48. Kim Y, Kang C, Min B et al. Detection and analysis of disease-associated single nucleotide polymorphism influencing post-translational modification, BMC Med Genomics 2015;8 Suppl 2:S7.

49. Wagih O, Reimand J, Bader GD. MIMP: predicting the impact of mutations on kinase-substrate phosphorylation, Nat Methods 2015;12:531–533.

50. Durek P, Schudoma C, Weckwerth W et al. Detection and characterization of 3D-signature phosphorylation site motifs and their contribution towards improved phosphorylation site prediction in proteins, BMC Bioinformatics 2009;10:117.

51. Pan Y, Wang Z, Zhan W et al. Computational identification of binding energy hot spots in protein-RNA complexes using an ensemble approach, Bioinformatics 2018;34:1473–1480.

52. Song J, Li F, Takemoto K et al. PREvaIL, an integrative approach for inferring catalytic residues using sequence, structural, and network features in a machine-learning framework, J Theor Biol 2018;443:125–137.

53. Li D, Mei H, Shen Y et al. ECharts: A declarative framework for rapid construction of web-based visualization, Visual Informatics 2018.

54. Rego N, Koes D. 3Dmol.js: molecular visualization with WebGL, Bioinformatics 2015;31:1322–1324.

55. Oates ME, Romero P, Ishida T et al. D(2)P(2): database of disordered protein predictions, Nucleic Acids Res 2013;41:D508–516.

56. Gille C, Birgit W, Gille A. Sequence alignment visualization in HTML5 without Java, Bioinformatics 2014;30:121–122.

57. Christensen B, Nielsen MS, Haselmann KF et al. Post-translationally modified residues of native human osteopontin are located in clusters: identification of 36 phosphorylation and five O-glycosylation sites and their biological implications, Biochem J 2005;390:285-292.

58. Brooks CL, Gu W. Ubiquitination, phosphorylation and acetylation: the molecular basis for p53 regulation, Curr Opin Cell Biol 2003;15:164-171.

59. Darling AL, Uversky VN. Intrinsic Disorder and Posttranslational Modifications: The Darker Side of the Biological Dark Matter, Front Genet 2018;9:158.

