## Supplementary material for "PRISMOID: a comprehensive 3D structure database for post-translational modifications and mutations with functional impact": The statistics of each type of PTM data are shown in Table S1

**Table S1.** List of top 100 ranked disease mutations in the PRISMOID database according to the number of mutations.

| **Disease Type** | **Num. of mutations** |
| --- | --- |
| Lissencephaly 3 | 80 |
| Cystic fibrosis | 67 |
| Lynch syndrome | 62 |
| Rasopathy, Noonan syndrome | 52 |
| Hereditary cancer-predisposing syndrome | 51 |
| Tumor predisposition syndrome | 48 |
| Rasopathy, Noonan syndrome, Noonan syndrome 1 | 36 |
| Noonan syndrome | 26 |
| Argininosuccinate lyase deficiency | 24 |
| Li-Fraumeni syndrome | 23 |
| Rasopathy, Noonan syndrome 1, Noonan syndrome | 22 |
| Costello syndrome | 22 |
| Familial hypercholesterolemia | 20 |
| Nephrotic syndrome, type 8 | 20 |
| Li-Fraumeni syndrome, Hereditary cancer-predisposing syndrome | 20 |
| Baraitser-Winter Syndrome 2 | 18 |
| Maple syrup urine disease | 17 |
| Li-Fraumeni syndrome 1, Li-Fraumeni syndrome, Tumor predisposition syndrome, Hereditary cancer-predisposing syndrome | 16 |
| Aicardi Goutieres syndrome 4 | 14 |
| Ornithine aminotransferase deficiency | 13 |
| Rasopathy, Noonan syndrome 1, Noonan syndrome, Early T cell progenitor acute lymphoblastic leukemia | 12 |
| Deafness, autosomal dominant 20 | 12 |
| Amyotrophic lateral sclerosis type 1 | 12 |
| Phosphoglycerate kinase 1 deficiency | 12 |
| Li-Fraumeni syndrome, Tumor predisposition syndrome, Hereditary cancer-predisposing syndrome | 12 |
| Juvenile myelomonocytic leukemia, Rasopathy | 12 |
| Tumor predisposition syndrome, Lynch syndrome | 11 |
| PI NULL(MATTAWA) | 11 |
| Hemorrhagic disease due to alpha-1-antitrypsin Pittsburgh mutation, Alpha-1-antitrypsin deficiency | 11 |
| PI CHRISTCHURCH | 11 |
| Hereditary cancer-predisposing syndrome, Lynch syndrome | 10 |
| Rasopathy | 10 |
| Epidermal nevus syndrome | 9 |
| Rasopathy, Noonan syndrome 1 | 8 |
| Parkinson disease 7 | 8 |
| Li-Fraumeni syndrome 1, Li-Fraumeni syndrome, Hereditary cancer-predisposing syndrome | 8 |
| Lynch syndrome, Hereditary cancer-predisposing syndrome | 8 |
| Non-small cell lung cancer | 8 |
| LEOPARD syndrome 1, Noonan syndrome, Rasopathy | 8 |
| Li-Fraumeni syndrome, Tumor predisposition syndrome | 8 |
| APOE2 VARIANT | 8 |
| Hyperinsulinism-hyperammonemia syndrome | 8 |
| Triosephosphate isomerase deficiency | 8 |
| Li-Fraumeni syndrome, Hereditary cancer-predisposing syndrome, Glioma susceptibility 1 | 8 |
| Familial type 3 hyperlipoproteinemia | 8 |
| Platelet glycoprotein IV deficiency | 7 |
| Purine-nucleoside phosphorylase deficiency | 7 |
| Leukemia, Philadelphia chromosome-positive, resistant to imatinib | 7 |
| Congenital disorder of glycosylation type 1t | 7 |
| Fibrosis of extraocular muscles, congenital, 3a, with or without extraocular involvement | 7 |
| Hyperthyroxinemia, familial dysalbuminemic | 6 |
| Mandibuloacral dysplasia with type B lipodystrophy | 6 |
| Microcephaly-capillary malformation syndrome | 6 |
| ALBUMIN CASERTA | 6 |
| Hemolytic anemia | 6 |
| Parkinson disease 13 | 6 |
| HEMOGLOBIN MAPUTO, HEMOGLOBIN G (COPENHAGEN) | 6 |
| Alloalbuminemia | 6 |
| Noonan syndrome 1, Noonan syndrome | 6 |
| HEMOGLOBIN ARYA | 6 |
| HEMOGLOBIN BELFAST, HEMOGLOBIN RANDWICK | 6 |
| HEMOGLOBIN DENVER, HEMOGLOBIN ILMENAU, HEMOGLOBIN MEQUON | 6 |
| Primary familial hypertrophic cardiomyopathy | 6 |
| HEMOGLOBIN HOWICK, HEMOGLOBIN ROTHSCHILD | 6 |
| HEMOGLOBIN ARTA, HEMOGLOBIN CHEVERLY | 6 |
| Deafness, autosomal recessive 1A, Non-syndromic genetic deafness, Hearing impairment | 6 |
| HEMOGLOBIN KURDISTAN, HEMOGLOBIN SINAI, HEMOGLOBIN L (FERRARA), HEMOGLOBIN HASHARON, HEMOGLOBIN SEALY | 6 |
| Deafness, autosomal recessive 1A, Deafness, autosomal dominant 3a, Non-syndromic genetic deafness, Hearing impairment | 6 |
| Inclusion body myopathy with early-onset paget disease and frontotemporal dementia | 6 |
| Non-small cell lung cancer, Medullary thyroid carcinoma, Noonan syndrome | 6 |
| 46,XY sex reversal 8 | 6 |
| Deafness, autosomal dominant 3a | 6 |
| HEMOGLOBIN HAMMERSMITH, HEMOGLOBIN CHIBA, Heinz body anemia | 6 |
| HEMOGLOBIN GAVELLO, HEMOGLOBIN AVICENNA | 6 |
| Homocystinuria, pyridoxine-nonresponsive | 6 |
| 2-methyl-3-hydroxybutyric aciduria | 6 |
| Adenine phosphoribosyltransferase deficiency | 6 |
| Cystathioninuria | 6 |
| alpha Thalassemia, beta^0^ Thalassemia | 6 |
| Medium-chain acyl-coenzyme A dehydrogenase deficiency | 5 |
| Baraitser-Winter syndrome 1 | 5 |
| Long QT syndrome | 5 |
| Hereditary cutaneous melanoma | 5 |
| Phosphoribosylpyrophosphate synthetase superactivity | 5 |
| Carbohydrate-deficient glycoprotein syndrome type I | 5 |
| Deafness, autosomal recessive 1A | 5 |
| Cystic fibrosis, ivacaftor response - Efficacy | 5 |
| HEMOGLOBIN MANAWATU | 4 |
| Arrhythmogenic right ventricular cardiomyopathy | 4 |
| Acute intermittent porphyria | 4 |
| HEMOGLOBIN ATHENS-GEORGIA | 4 |
| HEMOGLOBIN G (GALVESTON), HEMOGLOBIN G (TEXAS), HEMOGLOBIN G (PORT ARTHUR) | 4 |
| Lesch-Nyhan syndrome | 4 |
| beta Thalassemia | 4 |
| Amyotrophic lateral sclerosis 1, autosomal recessive | 4 |
| HEMOGLOBIN BOURMEDES | 4 |
| HEMOGLOBIN POITIERS | 4 |
| HEMOGLOBIN OITA, HEMOGLOBIN FORT DE FRANCE | 4 |
| HEMOGLOBIN AUSTIN | 4 |
| HEMOGLOBIN HANDSWORTH | 4 |
