## Supplementary material for "PRISMOID: a comprehensive 3D structure database for post-translational modifications and mutations with functional impact": Table 2 and Supplementary Table S1 list the cancer types and disease types of somatic cancer mutations and disease mutations, respectively.

**Table 1.** The statistics of each PTM type in the PRISMOID database.

| **PTM type** | **Target amino acid** | **Number of**  **Proteins** | **Number of PTM sites** | **URL address** |
| --- | --- | --- | --- | --- |
| Succinylation | Lysine | 1123 | 3578 | [Succinylation_K](http://prism.erc.monash.edu/pdb/browseType?type=Succinylation_K) |
| Ubiquitylation | Lysine | 1050 | 3540 | [Ubiquitylation_K](http://prism.erc.monash.edu/pdb/browseType?type=Ubiquitylation_K) |
| Phosphorylation | Serine | 1131 | 1996 | [Phosphorylation_S](http://prism.erc.monash.edu/pdb/browseType?type=Phosphorylation_S) |
| Phosphorylation | Tyrosine | 726 | 1238 | [Phosphorylation_Y](http://prism.erc.monash.edu/pdb/browseType?type=Phosphorylation_Y) |
| Acetylation | Lysine | 588 | 1212 | [Acetylation_K](http://prism.erc.monash.edu/pdb/browseType?type=Acetylation_K) |
| Glycation | Lysine | 384 | 1133 | [Glycation_K](http://prism.erc.monash.edu/pdb/browseType?type=Glycation_K) |
| N-linked Glycosylation | Asparagine | 534 | 1129 | [N-linked Glycosylation_N](http://prism.erc.monash.edu/pdb/browseType?type=N-linked%20Glycosylation_N) |
| Malonylation | Lysine | 514 | 1104 | [Malonylation_K](http://prism.erc.monash.edu/pdb/browseType?type=Malonylation_K) |
| Phosphorylation | Threonine | 702 | 1078 | [Phosphorylation_T](http://prism.erc.monash.edu/pdb/browseType?type=Phosphorylation_T) |
| Nitrosylation | Cysteine | 218 | 323 | [Nitrosylation_C](http://prism.erc.monash.edu/pdb/browseType?type=Nitrosylation_C) |
| Formylation | Lysine | 47 | 96 | [Formylation_K](http://prism.erc.monash.edu/pdb/browseType?type=Formylation_K) |
| Propionylation | Lysine | 47 | 90 | [Propionylation_K](http://prism.erc.monash.edu/pdb/browseType?type=Propionylation_K) |
| Methylation | Lysine | 45 | 82 | [Methylation_K](http://prism.erc.monash.edu/pdb/browseType?type=Methylation_K) |
| Acetylation | Serine | 65 | 67 | [Acetylation_S](http://prism.erc.monash.edu/pdb/browseType?type=Acetylation_S) |
| Acetylation | Alanine | 65 | 65 | [Acetylation_A](http://prism.erc.monash.edu/pdb/browseType?type=Acetylation_A) |
| O-linked Glycosylation | Serine | 32 | 64 | [O-linked Glycosylation_S](http://prism.erc.monash.edu/pdb/browseType?type=O-linked%20Glycosylation_S) |
| Sumoylation | Lysine | 45 | 61 | [Sumoylation_K](http://prism.erc.monash.edu/pdb/browseType?type=Sumoylation_K) |
| Glutarylation | Lysine | 17 | 46 | [Glutarylation_K](http://prism.erc.monash.edu/pdb/browseType?type=Glutarylation_K) |
| Acetylation | Methionine | 38 | 38 | [Acetylation_M](http://prism.erc.monash.edu/pdb/browseType?type=Acetylation_M) |
| O-linked Glycosylation | Threonine | 24 | 37 | [O-linked Glycosylation_T](http://prism.erc.monash.edu/pdb/browseType?type=O-linked%20Glycosylation_T) |
| Phosphoglycerylation | Lysine | 12 | 27 | [Phosphoglycerylation_K](http://prism.erc.monash.edu/pdb/browseType?type=Phosphoglycerylation_K) |
| Methylation | Arginine | 15 | 18 | [Methylation_R](http://prism.erc.monash.edu/pdb/browseType?type=Methylation_R) |
| Acetylation | Threonine | 17 | 17 | [Acetylation_T](http://prism.erc.monash.edu/pdb/browseType?type=Acetylation_T) |
| Crotonylation | Lysine | 11 | 17 | [Crotonylation_K](http://prism.erc.monash.edu/pdb/browseType?type=Crotonylation_K) |
| Palmitoylation | Cysteine | 11 | 16 | [Palmitoylation_C](http://prism.erc.monash.edu/pdb/browseType?type=Palmitoylation_C) |
| C-linked Glycosylation | Tryptophan | 5 | 16 | [C-linked Glycosylation_W](http://prism.erc.monash.edu/pdb/browseType?type=C-linked%20Glycosylation_W) |
| Carboxylation | Lysine | 11 | 11 | [Carboxylation_K](http://prism.erc.monash.edu/pdb/browseType?type=Carboxylation_K) |
| Acetylation | Glycine | 11 | 11 | [Acetylation_G](http://prism.erc.monash.edu/pdb/browseType?type=Acetylation_G) |
| Myristoylation | Glycine | 8 | 8 | [Myristoylation_G](http://prism.erc.monash.edu/pdb/browseType?type=Myristoylation_G) |
| Butyrylation | Lysine | 5 | 6 | [Butyrylation_K](http://prism.erc.monash.edu/pdb/browseType?type=Butyrylation_K) |
| 2-hydroxyisobutyrylation | Lysine | 3 | 5 | [2-hydroxyisobutyrylation_K](http://prism.erc.monash.edu/pdb/browseType?type=2-hydroxyisobutyrylation_K) |
| Lipoylation | Lysine | 4 | 4 | [Lipoylation_K](http://prism.erc.monash.edu/pdb/browseType?type=Lipoylation_K) |
| Prenylation | Cysteine | 4 | 4 | [Prenylation_C](http://prism.erc.monash.edu/pdb/browseType?type=Prenylation_C) |
| Carboxylation | Glutamic | 3 | 4 | [Carboxylation_E](http://prism.erc.monash.edu/pdb/browseType?type=Carboxylation_E) |
| Sulfation | Tyrosine | 2 | 2 | [Sulfation_Y](http://prism.erc.monash.edu/pdb/browseType?type=Sulfation_Y) |
| Neddylation | Lysine | 1 | 1 | [Neddylation_K](http://prism.erc.monash.edu/pdb/browseType?type=Neddylation_K) |
| Hydroxylation | Lysine | 1 | 1 | [Hydroxylation_K](http://prism.erc.monash.edu/pdb/browseType?type=Hydroxylation_K) |
