## Supplementary material for "PRISMOID: a comprehensive 3D structure database for post-translational modifications and mutations with functional impact": Table 2 and Supplementary Table S1 list the cancer types and disease types of somatic cancer mutations and disease mutations, respectively.

**Table 2.** Statistical summary of cancer mutations in the PRISMOID database.

| **Study name of the cancer type** | **Study Abbreviation** | **Num. of mutations** |
| --- | --- | --- |
| Bladder Carcinoma | BLCA | 1301 |
| Lung Squamous Cell Carcinoma | LUSC | 1155 |
| Lung Adenocarcinoma | LUAD | 1077 |
| Stomach Adenocarcinoma | STAD | 961 |
| Skin Cutaneous Melanoma | SKCM | 878 |
| Uterine Corpus Endometrial Carcinoma | UCEC | 832 |
| Head-Neck Squamous Cell Carcinoma | HNSC | 782 |
| Breast Invasive Carcinoma | BRCA | 698 |
| Colon Adenocarcinoma | COAD | 615 |
| Cervical Squamous Cell Carcinoma and Endocervical Adenocarcinoma | CESC | 532 |
| Liver Hepatocellular Carcinoma | LIHC | 433 |
| Esophageal Carcinoma | ESCA | 293 |
| Kidney Renal Clear Cell Carcinoma | KIRC | 279 |
| Low Grade Glioma | LGG | 268 |
| Prostate Adenocarcinoma | PRAD | 265 |
| Glioblastoma Multiforme | GBM | 228 |
| Cervical Kidney renal papillary cell carcinoma | KIRP | 213 |
| Sarcoma | SARC | 170 |
| Thyroid Cancer | THCA | 118 |
| Rectum Adenocarcinoma | READ | 88 |
| Pancreatic adenocarcinoma | PAAD | 82 |
| Diffuse Large B Cell Lymphoma | DLBC | 81 |
| Uterine Carcinosarcoma | UCS | 68 |
| Ovarian | OV | 67 |
| Adenoid cystic carcinoma | ACC | 58 |
| Kidney Chromophobe | KICH | 50 |
| Testicular germ cell tumor | TGCT | 32 |
| Uveal Melanoma | UVM | 24 |
| Mesothelioma | MESO | 24 |
| Pheochromocytoma and Paraganglioma | PCPG | 21 |
| Cholangiocarcinoma | CHOL | 16 |
| Thymoma | THYM | 13 |
